# Effects of Inbreeding on Microbial Community Diversity of *Zea mays*

**DOI:** 10.1101/2023.01.19.524730

**Authors:** Corey R Schultz, Matthew Johnson, Jason G Wallace

**Affiliations:** Institute of Bioinformatics, University of Georgia; Plant Breeding, Genetics, and Genomics, University of Georgia; Crop and Soil Science, University of Georgia

**Author notes:** Corresponding Author: Corey Schultz.

**Keywords:** Maize, microbiome, inbreeding, heterosis, genetic variation

## Abstract

Heterosis, also known as hybrid vigor, is the basis of modern maize production. The effect of heterosis on maize phenotypes has been studied for decades, but its effect on the maize-associated microbiome is much less characterized. To determine the effect of heterosis on the maize microbiome, we sequenced and compared the bacterial communities of inbred, open pollinated, and hybrid maize. Samples covered three tissue types (Stalk, Root, and Rhizosphere) in two field experiments and one greenhouse experiment. Bacterial diversity was affected by location and tissue type, but not genetic background, for both within-sample (alpha) and between-sample (beta) diversity. PERMANOVA analysis similarly showed that tissue type and location had significant effects on the overall community structure, whereas the genetic background and individual plant genotypes did not. Differential abundance analysis identified only 18 bacterial ASVs that significantly differed between inbred and hybrid maize. Predicted metagenome content was inferred with Picrust2, and it also showed a significantly larger effect of tissue and location than genetic background. Overall, these results indicate that the bacterial communities of inbred and hybrid maize are often more similar than they are different, and that non-genetic effects are generally the largest influences on the maize microbiome.

## 1. Introduction

All plants coexist with communities of fungi and bacteria in, on, and around them [1, 2]. These microbes can colonize aboveground surfaces (the phyllosphere), soil near the roots (the rhizosphere), and the interior of plant tissues (the endosphere) [1, 3], and they can significantly contribute to the overall health of the plant [1]. A plant’s microbiota—the collection of all microbes associated with it—can benefit the plant by protecting it from pathogens and herbivores [4–6], protecting against abiotic stress [7–11], and promoting growth through nutrient acquisition (through N fixation, P solubilization, siderophore production, etc) [2, 12–16], and phytohormone production [17–19]. Beneficial endophytes can activate plant immune responses resulting in a level of protection from pathogens [20–22]. In turn, the host plants affect microbes by changing soil chemistry and secreting signaling compounds [23–26], exuding energy-rich carbon compounds into the rhizosphere [24], and otherwise providing niches for microbes [27].

An active research area in plant-microbe interactions is determining the extent to which plant genetic variation alters the microbial community [2, 28–30]. One motivation for this research is the idea of breeding crops for improved microbial associations [28]. Several studies have shown that host genetics significantly affects microbiome community structure in maize [3, 31, 32], rice [33, 34], wheat [35, 36], and other crops [37–39].

Maize has been a model crop for plant genetics for over 100 years [40], due in a large part to its extensive genetic variation and high economic value (170.7 bushels/acre and $9.2 billion in US exports alone in 2020) [41–43]. Although most commercial maize consists of F1 hybrids [41], most maize microbiome research has been on inbred lines [3, 31, 44, 45], with only a few studies examining the difference between inbred and hybrid maize [32, 46].

F1 hybrids show increased vigor and yield relative to their parents [32], an effect called hybrid vigor or heterosis. Heterosis, which is particularly strong in maize, can manifest as increased growth rate, biomass, stress resistance, and yield [47, 48].

Recently it was found that field-grown maize displays heterosis in bacterial rhizosphere communities, as well as fungal communities in the rhizosphere and phyllosphere [32]. In addition, heterosis for germination and root biomass was shown, at least in some instances, to depend upon the local microbial community [46]. In this case, heterosis resulted from inbreds performing as well as hybrids in sterile conditions but worse in the presence of microbes. These results indicate that some part of heterosis may be due to hybrids’ superior ability to deal with harmful microbes in the environment.

These previous studies focused on the exterior communities of the plant (the rhizosphere and phyllosphere). In this study we sought to characterize how inbreeding and heterosis affects both the interior and exterior bacterial communities of maize by looking at the bacteria of the rhizosphere, root endosphere, and stalk endosphere communities in three differently inbred maize groups (inbreds, F1 hybrids, and open-pollinated varieties). Our primary goals were to (1) characterize the bacterial communities in each compartment for each group, (2) determine aspects of the community that were consistent across them, (3) determine differences in the communities that could be linked to heterosis, and (4) test the hypothesis that hybrid maize may be selecting superior microbial communities.

## 2. Materials and Methods

### Maize Cultivars

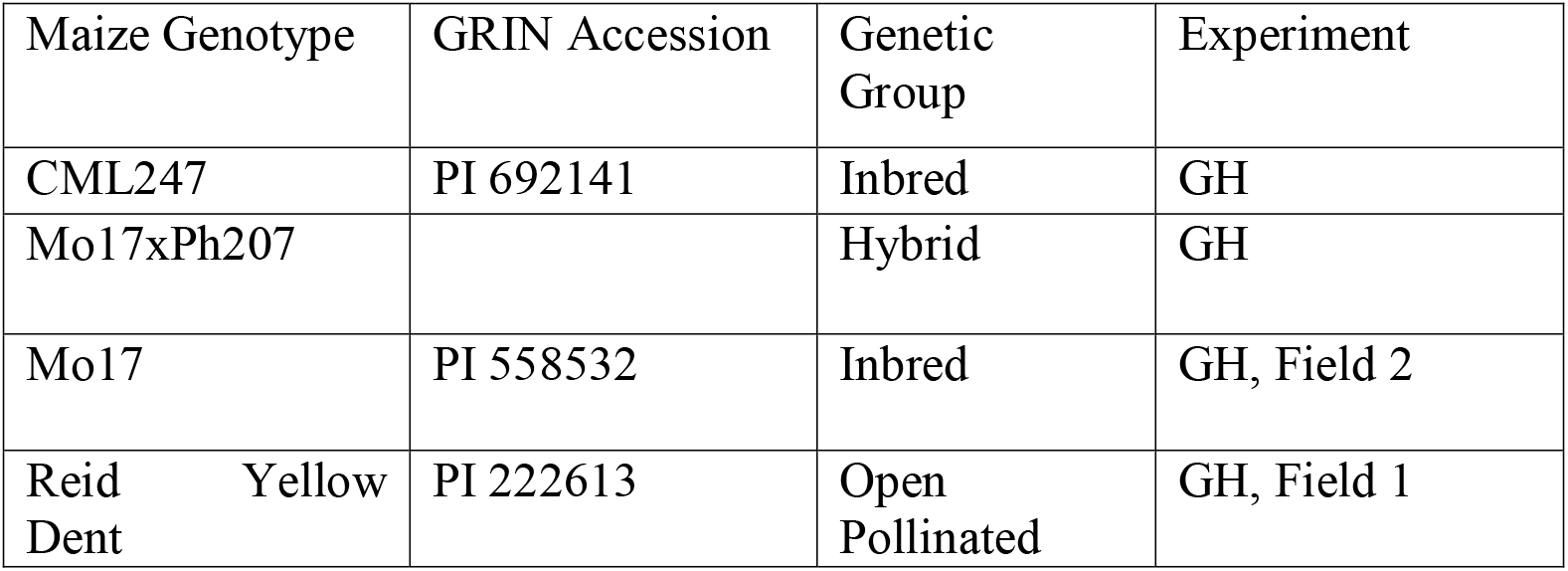
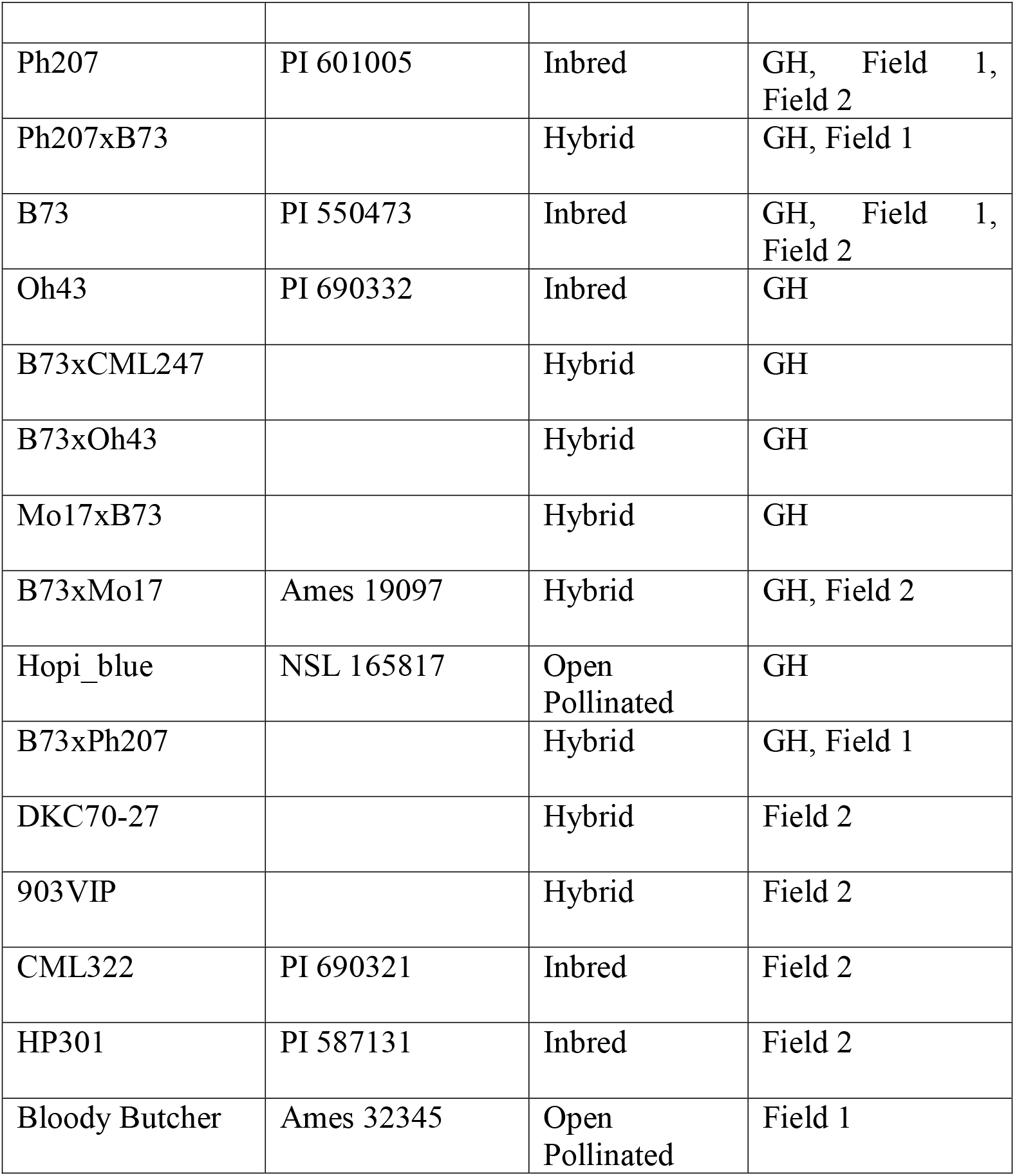

| Maize Genotype | GRIN Accession | Genetic Group | Experiment |
| --- | --- | --- | --- |
| CML247 | PI 692141 | Inbred | GH |
| Mo17xPh207 |  | Hybrid | GH |
| Mo17 | PI 558532 | Inbred | GH, Field 2 |
| Reid Yellow Dent | PI 222613 | Open Pollinated | GH, Field 1 |
| Ph207 | PI 601005 | Inbred | GH, Field 1, Field 2 |
| Ph207xB73 |  | Hybrid | GH, Field 1 |
| B73 | PI 550473 | Inbred | GH, Field 1, Field 2 |
| Oh43 | PI 690332 | Inbred | GH |
| B73xCML247 |  | Hybrid | GH |
| B73xOh43 |  | Hybrid | GH |
| Mo17xB73 |  | Hybrid | GH |
| B73xMo17 | Ames 19097 | Hybrid | GH, Field 2 |
| Hopi_blue | NSL 165817 | Open Pollinated | GH |
| B73xPh207 |  | Hybrid | GH, Field 1 |
| DKC70-27 |  | Hybrid | Field 2 |
| 903VIP |  | Hybrid | Field 2 |
| CML322 | PI 690321 | Inbred | Field 2 |
| HP301 | PI 587131 | Inbred | Field 2 |
| Bloody Butcher | Ames 32345 | Open Pollinated | Field 1 |

### Field and Greenhouse Design

A single greenhouse experiment was carried out in 2019. For each pot, four seeds were planted in a 5 gallon pot with 90% Professional Growing Mix Fafard 3B/Metro-Mix 830 (Sungro Horticulture) and 10% Vermiculite. Upon emergence pots were thinned down to one plant per pot. Pots were arranged in a randomized block design with three replicates, with each table in the greenhouse consisting of a block.

Fields were planted in the summer of 2018 and 2019 at the Iron Horse Research Farm in Watkinsville, Georgia. Plants were grown via standard agronomic practices for the state of Georgia [49].

### Sample Collection and Processing

Plants were harvested in a single day to avoid batch effects. A 10cm section of stalk was cut from the plant 20cm off the ground using sterilized razor blades and gloves. Plants were dug up around the roots, and roots were removed from the center of the root ball and placed into a clean falcon tube for root and rhizosphere samples.

The outer portion of stalks were removed with a sterile razor blade, and the inner tissue (protected from contamination and external microbes) were cut into 1-3mm pieces and loaded into a 2mL conical tube for GenoGrinding (SPEX SamplePrep). Root samples were vortexed on the max setting for 15s in deionized water to separate the rhizosphere from the root. This wash was then centrifuged at 4500g for 10 minutes to prepare the wash for DNA extraction. Roots were then thoroughly cleaned with deionized water to remove any residual rhizosphere. 2-3cm of roots were chopped up with a sterile razor blade and loaded into a 2mL conical tube for GenoGrinding (SPEX SamplePrep).

### DNA Extraction and Sequencing

DNA was extracted with a Quick-DNA Fecal/Soil Microbe 96 Kit (Zymo), following the manufacturer’s instructions. 16s rDNA gene amplification was performed using the Earth Microbiome Project 515F(Parada) and 806R(Apprill) primers (Thompson, et.al. 2017) with linkers. Sequences are GTGYCAGCMGCCGCGGTAAGT (515F) and GGACTACNVGGGTWTCTAATCC (806R). Peptide nucleic acids (pPNA and mPNA, to block plastid and mitochondrial amplification, respectively; PNA Bio) were mixed and diluted to 2.5uM each for inclusion in the reaction. The first PCR reaction consisted of 5 μL DNA template, 2 μL of each primer (0.5 μM), 12.5 μL of Hot Start Taq 2X Master Mix (New England Biolabs), 2.5 μL PNA mixture (2.5 μM each), and 1uL of sterile water. The amplification reaction was 95°C for 45 seconds; twenty cycles of 95°C for 15 seconds, 78°C for 10 seconds, 60°C for 45 seconds, 72°C for 45 seconds; and finally hold at 4°C. PCR products were purified with AMPure (Beckman Coulter Life sciences).

Five μL of the first PCR product for each sample was used in the second PCR amplification. The reaction mix consisted of 5 μL first PCR product, 5 μL Nextera i5 and i7 Barcode Primers, 12.5 μL 2x Taq DNA polymerase master mix, and 2.5 μL PNA mix. The second amplification reaction was 95°C for 45 seconds; 25 cycles of 95°C for 15 seconds, 78°C for 10 seconds, 60°C for 45 seconds, 72°C for 45 seconds; and finally, 68°C for 5 minutes followed by hold at 4°C. The second PCR products were purified using AMPure beads and the cleaned products were eluted in 27uL of sterile water and stored at −20°C until sequencing. Three blanks were used in DNA extraction and library prep. Libraries were sequenced at the Georgia Genomics and Bioinformatics Core on an Illumina MiSeq instrument using one paired-end 250 flowcell. The raw data are available at the NCBI Sequence Read Archive under accession PRJNA924784.

### Bioinformatics

Sequence processing and quality filtering was completed within the Qiime2 version 2019.1 toolbox [50]. Cutadapt [51] was used to trim primers from raw sequences and filter reads that did not reach a Phred score of 26. FastQC was used to visualize read quality [52]. Paired reads were joined with vsearch in Qiime2 [53]. deblur [54] was used to truncate reads to 200 bp. The SILVA 132-99-nb classifier [55] was used to assign taxonomy to ASVs. The original dataset contained 15126 ASVs in 252 samples. Taxa with no Phylum identity were discarded, as well as ASVs found in blanks, and ambiguous calls. We also removed taxa related to the host, chloroplasts, and mitochondria, as well as ASVs that were not present in at least 5% of samples. ASVs were not agglomerated into OTUs. This left us with 4628 ASVs. For some analysis (like Core ASVs) we only looked at the 738 taxa that were found in all three experiments.

First we compared alpha diversity based on genetic background, tissue, and location. We used Observed ASVs, Shannon, and Simpson index from the phyloseq package [56]. Pairwise Wilcox tests and Dunn’s post-hoc test were used to test for significance with the FSA package [57]. UniFrac distance matrices were generated in qiime2 for beta diversity and plotted to visually represent sample diversity. To test what variables had the most significant impact on beta diversity, we generated Bray-Curtis distances in phyloseq, and then a PERMANOVA was performed using vegan [58]. Differential abundance analysis was used to identify ASVs that occurred in all three experiments that differed between inbred and hybrid maize. DESeq2 [59] was used to fit negative binomial models with an alpha value of 0.001. Core ASVs were defined as present in 90% of samples. UpSet plots were created using the UpSetR [60] package. PiCRUST2 [61] was used to predict functional gene pathways from ASVs using the Kegg Orthology database [62]. Raw KO terms were agglomerated to higher functional pathways, and DESeq2 was used to identify pathways that differed between inbred and hybrid maize compartments, with an alpha value of 0.001. psadd was used to create interactive krona plots of microbiome taxonomy [63, 64]. All bioinformatics scripts and pipelines are available at: https://github.com/wallacelab/paper-schultz-microbiome-2023

### MiniMaize Inoculation Experiment

Maize lines B73, Mo17, and their F1 hybrid seeds were sterilized via our previously established method [65]. Seeds were surface sterilized with sterile water, bleach, and tween 20 and then placed in a hot water bath. They were then allowed to germinate on Hoagland’s agar for seven days to check for contamination. These seeds were then planted in the greenhouse as described above and grown until flowering. Once the silks emerged, stalk sections were sampled from 6 – 12 inches above the soil line. A razor was used to cut a 10 cm x 10 cm square in the side of mature maize root ball, and roots were removed from the plant to include roots all the way to the center of the pot.

Microbiome extraction was modified from [66, 67]. Stalk samples were cored using a sterilized drill tip. Stalk pulp was placed into a 50mL falcon tube, and filled with 40mLs of MilliQ H2O, then shaken 50 times and vortexed on max speed for 10 seconds. 30mLs of liquid was decanted into another falcon tube, using the tube cap to exclude large debris. The microbe suspension was centrifuged for 2 minutes at 3,500 RPM to pellet plant debris. 20 mLs of this was filtered through a 2 micrometer Whatman filter. Root samples were placed into a falcon tube without drill tip pulverization, and the same method was used to extract rhizosphere microbiomes.

Sterilized MiniMaize seeds were planted in 2.7L sterilized pots, with autoclave media mixture (same as above). Two sterilized MiniMaize seeds were planted in each pot. Pots were inoculated with 10 mLs of either B73, Mo17, or F1 combined stalk and rhizosphere microbiomes, with 8 pots per treatment. Autoclaved tin foil was placed over the pots for 4 days to ensure no outside microbes were introduced, then plants were thinned to one plant per pot. Plants were allowed to grow for 5 weeks. Shoots were cut at the soil line and placed in brown paper bags. Roots were gently washed of soil and bagged. Above and below ground samples were dried out and weighed.

## 3. Results

We grew hybrid, inbred, and open pollinated maize lines in two field experiments (2018 and 2019) and one greenhouse experiment. Bacterial microbiomes were extracted from stalks, roots and rhizospheres of the plants, and quantified with QIIME2 and deblur. High-quality reads were retained and classified with SILVA taxonomy classifier. Low-abundance amplicon sequence variants (ASVs), as well as ASVs associated with mitochondria, chloroplasts, and our blanks, were filtered out. Our final dataset consisted of 252 samples and 2806 ASVs, 738 (26%) of which were present across all three experiments.

Throughout the three experiments we found that stalk tissue had lower read depth and fewer associated ASVs compared to the rhizosphere and roots. Rhizosphere had the most classified ASVs, and most of them were shared by root samples (Figure 1A). Very few bacteria were shared between stalks and the other two tissues, though these shared ASVs accounted for the majority (99%) of stalk reads. The relative abundance of ASVs in the rhizosphere and root relative was roughly in line with the number of unique ASVs. Stalk samples, however, were dominated by Proteobacteria reads (36.1% of stalk ASVs but 74.9% of total read depth). Krona plots (nested pie chart distributions) of overall community structures, with comparisons for tissue and genetic background, can be found in the Supplemental Materials Dataset S1.

**Figure 1.**
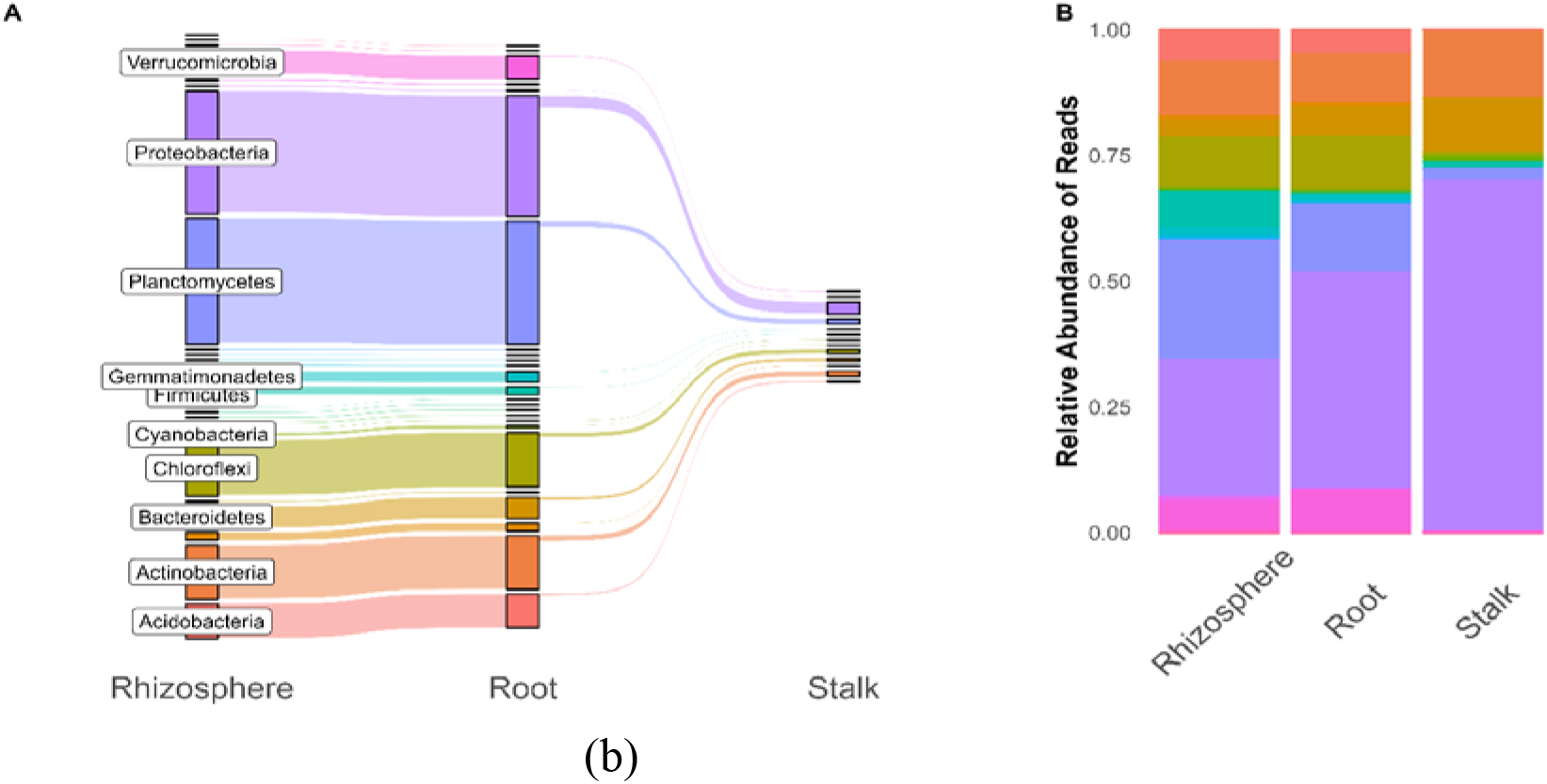
Shared ASVs across plant tissues, compared to relative abundance of reads in plant tissues, colored by plylum. The rhizosphere had the largest number of ASVs, with a majority of these also found in the roots. Only a fraction of the microbial community found in the rhizosphere and roots can be found in the stalks. While the number of ASVs in a phylum appear to accurately represent the relative abundance of reads in the rhizosphere and root, we see differences in the stalk. Although planctomycetes make up a sizable proportion of ASVs in the stalk, they represent a very small amount of reads, which is mainly dominated by Proteobacteria.

To investigate what taxa were shared by groups of samples, we plotted intersections of common ASVs collapsed at the genus level (Figure 2). We defined common taxa as genera that were found in at least 50% of samples in a group; a table for all taxa and groups can be found in the Supplemental Materials. Forty-four genera were shared by at least 50% of samples in inbred and hybrid roots and rhizospheres. 10 genera were shared by inbred roots and rhizospheres but not hybrids, and vice versa. As a group, the rhizosphere samples contained 5 genera that were not found in the roots, while the roots had 3 genera not found in the rhizosphere. Inbred and hybrid stalk samples only shared one common microbe that was not found in underground compartments. No genera were shared across all samples and genetic backgrounds. All intersections can be found is Table S1.

**Figure 2.**
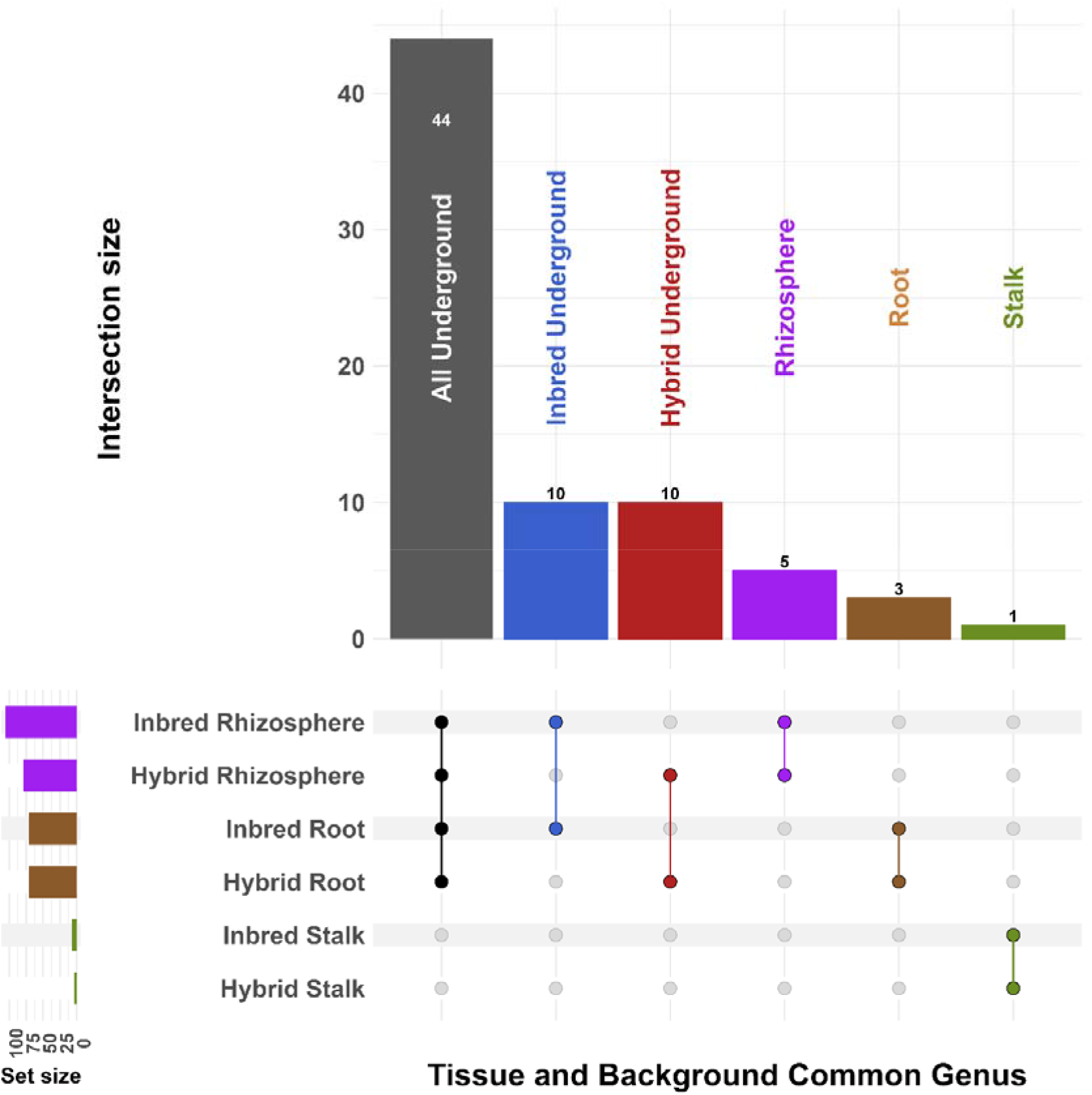
UpSet plot [60] showing intersections of common genera (those found in >50% of samples) based on genetic background and tissue. Intersections with zero counts are not shown. The largest number of shared genera are in the underground compartments and are shared across both inbred and hybrid material. Stalk samples, meanwhile, were much less diverse and contained a single shared genus between inbred and hybrid varieties. Open pollinated lines share genera in some, but not all of these intersections.

Alpha diversity was measured with three common metrics--Observed ASVs, Shannon Entropy, and Hill’s q1 [exponential of Shannon Entropy [68] on rarefied data (Figure S1)—and compared using Kruskal-Wallis and Dunn’s test. For all tissue types, we find that field samples have higher alpha diversity than their greenhouse counterparts (p < .001). Similarly, the root and rhizosphere samples have higher alpha diversity than stalk samples (p < .001). Post-hoc tests show that there are no significant differences in alpha diversity comparing inbred, hybrid, or open-pollinated samples across experiments (Table S2).

Beta diversity was calculated using the Weighted UniFrac metric [69] (Fig. 3). Samples were most strongly separated based on tissue type, with rhizosphere, root, and stalks strongly separating from each other. Whether the experiment made a difference depended on the tissue: rhizosphere and root samples were strongly differentiated based on experiment, and stalk samples not at all. Genetic background did not significantly differentiate samples in any compartment. PERMANOVA analysis of Weighted UniFrac distances indicated that experiment and tissue type had the most impact on beta diversity (p = .001 and p = .001 by Type II ANOVA) (Table S3). Genetic background (inbred/hybrid/open-pollinated) and individual genotype had no significant effect on beta diversity (p > 0.05; data not shown).

**Figure 3.**
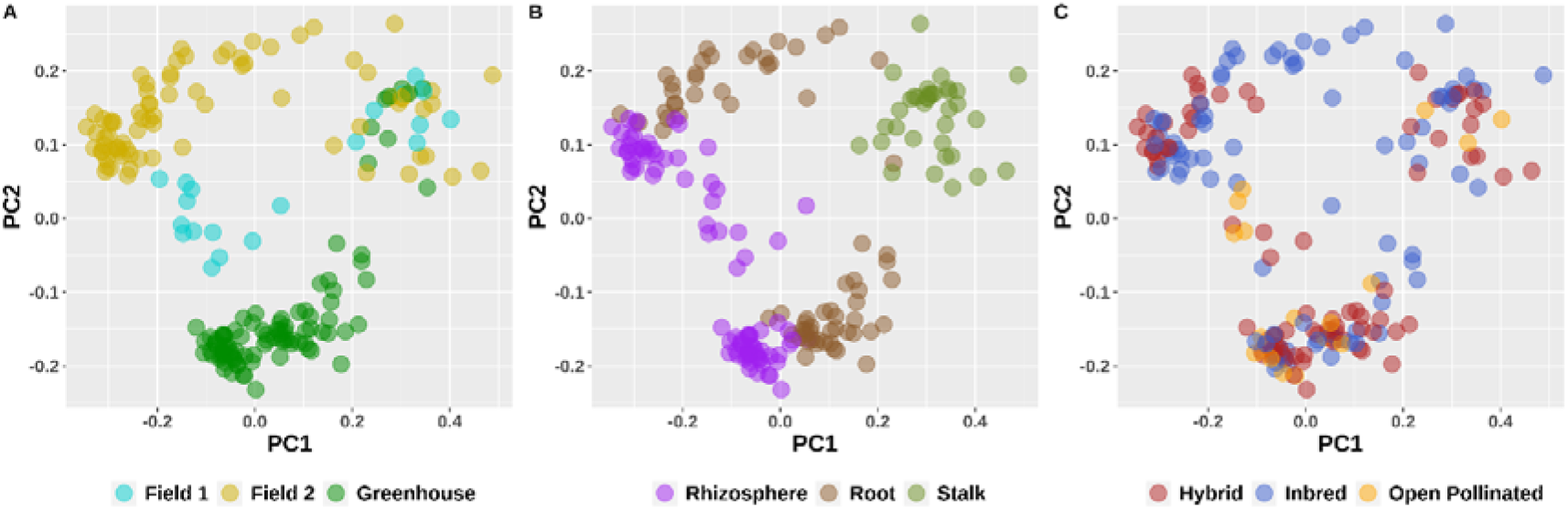
Weighted UniFrac diversity principle coordinates, colorized by experiment (A), tissue type (B), and genetic background (C). Samples separate by location and tissue type but not genetic background.

To identify the microbes that are most different between samples, we analyzed differentially abundant microbes using DESeq2 (Table 1 and Fig 4), using the 738 ASVs found in all three experiments. Table 1. shows differential comparisons across tissue type, genetic background, and location. We see that location and tissue type have far more differentially abundant microbes than comparisons of genetic background. Most of these were found in the root (11/18), and many of these were members of the Burkholderiaceae and Rhizobiaceae families.

**Table 1.** Table of Differentially abundant taxa. DESeq2 was used to compare genetic background, location, and tissue type. Tissue type and location had a much large impact on the number of differentially abundant ASVs than genetic background.

| **Tissue** | **Comparison** | **Number of ASVs** |
| --- | --- | --- |
| *Compare genetic background* |  |  |
| All | Inbred vs Hybrid | 55 |
| All | Inbred vs Open Pollinated | 78 |
| All | Hybrid vs Open Pollinated | 25 |
| Rhizos | Inbred vs Hybrid | 2 |
| Rhizos | Inbred vs Open Pollinated | 5 |
| Rhizos | Hybrid vs Open Pollinated | 6 |
| Roots | Inbred vs Hybrid | 5 |
| Roots | Inbred vs Open Pollinated | 11 |
| Roots | Hybrid vs Open Pollinated | 2 |
| Stalks | Inbred vs Hybrid | 5 |
| Stalks | Inbred vs Open Pollinated | 5 |
| Stalks | Hybrid vs Open Pollinated | 5 |
| *Compare locations* |  |  |
| All | Field vs Greenhouse | 475 |
| Rhizos | Field vs Greenhouse | 177 |
| Roots | Field vs Greenhouse | 188 |
| Stalks | Field vs Greenhouse | 24 |
| *Compare tissues* |  |  |
| - | Stalks vs Rhizos | 433 |
| - | Stalks vs Roots | 355 |
| - | Roots vs Rhizos | 275 |

**Figure 4.**
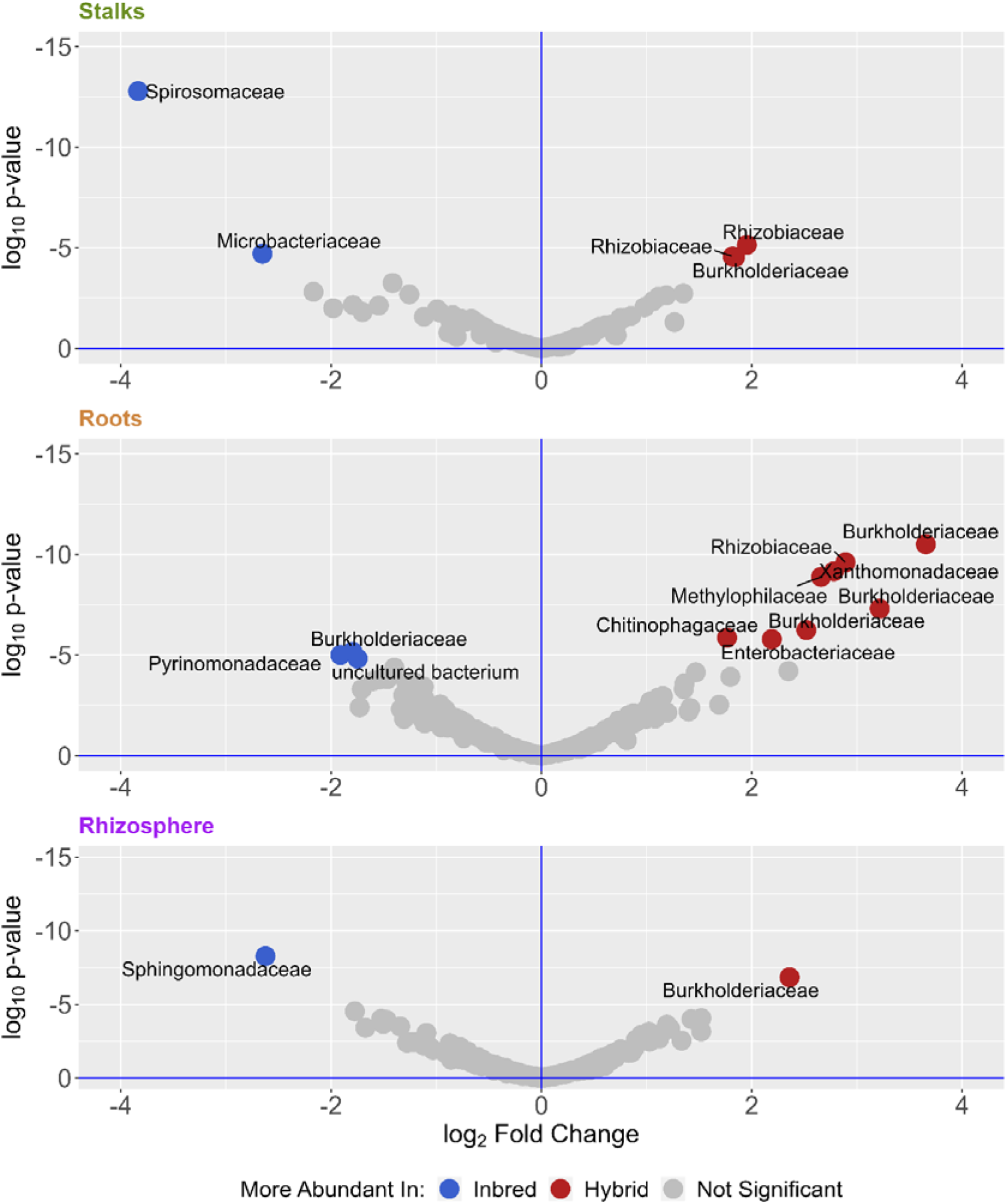
Volcano plots of differentially abundant ASVs. ASVs were more abundant in Inbred (blue) or hybrid (red) determined by DESeq2 with an alpha of .001. Roots had the most differentially abundant ASVs. Dot represent individual ASVs, which are labeled according to their taxonomic Family. The full list of differentially abundant ASVs is in Supplemental Table 4.

Previous work has shown that metabolic functions are a better characterization of microbial communities than 16s-based taxonomy [31, 70, 71].We used PICRUST2 [61] to predict community functional capacity from the 16s sequencing, and DESeq2 to compare differential abundance for comparisons across genetic background, tissue, and location (Table 2). Functional differences were much larger between tissue types and locations than genetic background. When comparing inbred and hybrid tissues, while each compartment had some individually different metabolic functions, only the roots maintained significant differences when individual functional annotations were grouped into metabolic pathways (Table 2). Inbred roots had an increase in predicted gene groups related to metabolism and molecular degradation, while hybrid maize roots had an increase in groups related to carbon transport, electron transfer carriers, and photosynthesis (Figure S2).

**Table 2.**
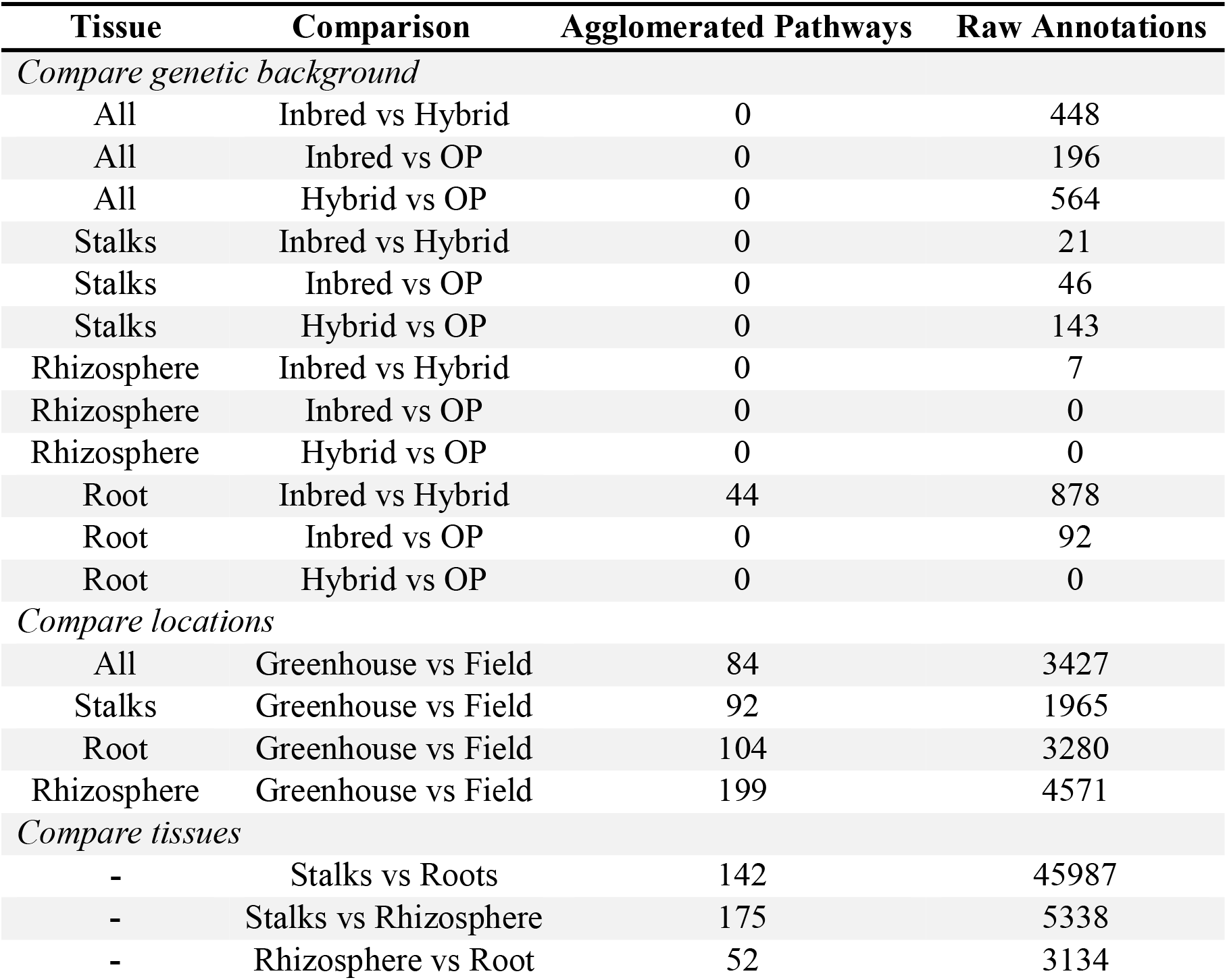
Differentially abundant PICRUST predicted genomic functions. Agglomerated pathways were grouped based on KEGG pathways before differential abundance was performed (alpha of .001). Comparisons between genetic backgrounds were much smaller compare

| **Tissue** | **Comparison** | **Agglomerated Pathways** | **Raw Annotations** |
| --- | --- | --- | --- |
| *Compare genetic background* |  |  |  |
| All | Inbred vs Hybrid | 0 | 448 |
| All | Inbred vs OP | 0 | 196 |
| All | Hybrid vs OP | 0 | 564 |
| Stalks | Inbred vs Hybrid | 0 | 21 |
| Stalks | Inbred vs OP | 0 | 46 |
| Stalks | Hybrid vs OP | 0 | 143 |
| Rhizosphere | Inbred vs Hybrid | 0 | 7 |
| Rhizosphere | Inbred vs OP | 0 | 0 |
| Rhizosphere | Hybrid vs OP | 0 | 0 |
| Root | Inbred vs Hybrid | 44 | 878 |
| Root | Inbred vs OP | 0 | 92 |
| Root | Hybrid vs OP | 0 | 0 |
| *Compare locations* |  |  |  |
| All | Greenhouse vs Field | 84 | 3427 |
| Stalks | Greenhouse vs Field | 92 | 1965 |
| Root | Greenhouse vs Field | 104 | 3280 |
| Rhizosphere | Greenhouse vs Field | 199 | 4571 |
| *Compare tissues* |  |  |  |
| - | Stalks vs Roots | 142 | 45987 |
| - | Stalks vs Rhizosphere | 175 | 5338 |
| - | Rhizosphere vs Root | 52 | 3134 |

Since hybrid maize is generally more fit than inbred maize, we hypothesized that hybrid maize would cultivate a more optimal microbiome. To test this, we grew the inbred lines B73 and Mo17, and their F1 hybrid to maturity from surface-sterilized seeds germinated in vitro. These plants were grown to flowering in the greenhouse, and were allowed to recruit microbiomes from non-sterile potting mix. Fast-Flowering Mini-Maize [72] was surface sterilized and germinated in vitro, then inoculated with filtered bacterial microbiomes harvested from stalks and rhizospheres of mature B73, Mo17, their F1 Hybrid, or an autoclaved control. After 28 days, roots and shoots were harvested, dried, and weighed. No significant differences occurred below-ground for the separate inoculation groups. Inoculation with microbiomes from B73 (p = 0 .048) and the F1 hybrid (p = 0.037) resulted in a small but significant decrease in shoot biomass relative to control (Figure S3). No other differences were significant.

## 4. Discussion

Our results show that inbreeding has a small but significant effect on maize microbial communities. These effects however, are much smaller than the effect that environment and tissue compartment have on microbiome makeup and predicted function. These results have several implications for maize-microbe interactions.

First, we showed that root communities were very similar to the rhizosphere soil, while only a fraction of these bacteria can be found in stalk tissue (Figure 1). This may be due to a combination of strong filters as bacteria travel up the plant [73, 74], or a larger effect of seed-transmitted microbes in the stalks compared to the roots [73]. Stalk samples had fewer ASVs overall and lower reads, and had a much smaller common microbiome when compared to root and rhizosphere (Figure 2). Although shared taxa were significantly different based on tissues, there were no such differences when comparing inbred and hybrid maize.

Similarly, there were no significant differences in alpha diversity measures based on inbreeding, but location and tissue had large affects (Figure S1). This is similar to the results of Wagner et al. [32], who also did not find significant differences in alpha diversity between inbred and hybrid maize. We found alpha diversity to be higher belowground than above ground, and we found field microbial communities to have higher alpha diversity than greenhouse communities. A similar pattern held for beta diversity, where tissue and experiment had larger impacts than inbreeding, as shown by both PCoA plots and PERMANOVA (Figure 3). These results add to the emerging collection of data that (1) maize tissues have different and distinct microbiomes [3, 31, 44], and (2) environment has a larger impact on plant microbiome assembly than genetics [3, 31, 75].

When directly comparing inbred versus hybrid communities, we identified 11 differentially abundant ASVs in the roots, 5 the stalks, and 2 in the rhizosphere (Figure 4). This represent a very small number of the ASVs in this dataset, and is much smaller than the hundreds of differential ASVs indicated by other characteristics (Table 1). Most of these differences were found in the roots, this may indicate the effect of inbreeding may be most pronounced there, or it may simply reflect that the root microbiomes were the largest and most diverse. Perhaps not surprisingly, many of these ASVs belong to groups known to include plant-associated and plant-beneficial bacteria, such as Rhizobiaceae and Burkholderiacea. Endophytes from both of these families have been shown to have growth promotion potential in maize [14, 76–78]. We also identified a species of the genus Pantoea, which is known to associate with plants [79–82], as well as a species of the genus Sphingomonas, which has been shown to promote growth and can play a role in phytoremediation [83].

Although predicting metabolic capacity from 16s data is less precise than actual metagenomics data, prior work has shown that metabolomics data supports PICRUST2 predictions [84]. It has been found that metabolic functions of the microbiome community may be more important than the taxonomic identity of the individual microbes [70, 71, 85], and Wallace et al [31], found predicted metabolic pathways to be more heritable (meaning affected by host genetics) than individual microbes in the maize leaf microbiome. In our data, we found predicted differences in gene functions in maize roots--but not the stalk or rhizosphere--when comparing inbred and hybrid maize. Similar to taxonomic differences (Table 1), the predicted metabolic differences were much smaller when comparing inbreeding than comparing locations and tissues (Table 2). Inbred roots had an increase in predicted gene groups related to acetyl-coA activity, molecular degradation of organic compounds, and increases in plant-microbe signaling; while hybrid maize roots had an increase in groups related to ribosome synthesis, energy, and photosynthetic functions (Figure S2). Taxa contributions to these gene groups (Dataset S3) were taxonomically diverse, and most were not found to be differentially abundant in inbred or hybrid roots, highlighting the importance of comparing community function. It has been shown that plant associated bacteria genomes encode more carbohydrate metabolism functions, as well diverse functions related to organic compound metabolism, and plant protein mimicry [86].

Inbred plants showed increases in acetyl-coA-transferases and dehydrogenases, implying there may be a difference in anaerobic metabolism, even though there were no known acetogens [87] in our differentially abundant ASVs. There were also increases in enzymes related to organic compounds that may be produced by the plant or bacteria. Plants produces exudes and can respond to stress through volatile organic compounds (VOCs), while plant microbiomes can also produce a wide array of VOCs that can impact plant health. These VOCs are incredibly varied and include alcohols, aldehydes, acids, esters, fatty acids, and hydrocarbons (reviewed in [88, 89]). We found predicted functions related to esters [88], hydrogen peroxide (plant VOC) ligases [90, 91], and phosphonates, which can protect bacteria and make phosphate available to the plant [92–94]. These increases may indicate differential communication between the plant and the microbiome, or the microbiome’s reaction to plant stress response. If heterosis is partly microbe-dependent [46], a thorough investigation of these signaling pathways may reveal mechanisms influencing heterosis in maize.

Hybrid plants had an increase in pathways related to ribosome biogenesis (including synthases and GTPase A), energy production, and photosynthesis/carbon metabolism. Several pathways related to energy production and storage were found increased in hybrid maize. These included oxidative phosphorylation, heme uptake proteins, and electron transport. Outside of these energy related pathways, a number of energy and protein pathways related to phototrophic capabilities were differentially abundant in hybrid maize root bacterial communities. These include: plastocyanin [95], cytochrome c subunit [95], and photo reaction center m [96, 97]; all related to photosynthetic energy production in photo-reactive bacteria [97]. Cyanobacteria ASVs, found within these maize microbiomes (Figure 1), had little effect on the differences in functional gene prediction. Differences in photosynthetic protein and photosynthesis gene groups were due to a wide range of phylums, including Proteobacteria, Chloroflexi, Fimicutes, and Acidobacteria (Dataset S3), all shown to have phototrophic capabilities [98]. These taxa were not identified as differentially abundant (Figure 4). This analysis indicates that hybrid plants may be selecting for microbial communities that have increased energy production or phototrophic potential. Actual metagenomic data, instead of 16s-based predictions would allow us the parse apart real metabolic differences between the two communities, whereas this analysis focuses on predicted potential of the communities.

When we inoculated MiniMaize with the filtered bacteria microbiome from B73, Mo17, and their F1 hybrid, we saw minor differences in biomass above ground but not below (Figure S3). Specifically, we saw B73 and F1 inoculates decreased plant biomass compared to control inoculated plants, while Mo17 had no significant effect. We hypothesized that hybrid maize may harbor more beneficial bacteria, but our results do not support this conclusion. In a previous study Kaeppler et al. [99] tested the mycorrhizal responsiveness of 28 inbred lines maize lines. It was found that B73 and Mo17 had the largest differences in response to mycorrhizae. Our study used bacterial filtrates, indicating these two lines may consistently respond differently to microbial inoculation.

Our results are somewhat similar with those of Wagner et al., who showed significant differences in inbred and hybrid maize microbial-communities in the field [32], and that at least some effects of heterosis may be due to hybrids’ superior ability to deal with harmful microbes [45].

Our primary goals were to (1) characterize the bacterial communities in each compartment for each group, (2) determine aspects of the community that were consistent across groups, (3) determine differences in the communities that could be linked to heterosis, and (4) test the hypothesis that hybrid maize may be selecting superior microbial communities.

We found that (1) bacterial communities in the roots and rhizospheres were very similar to each other, and stalk communities only have a small portion of these bacteria, (2) common bacteria were found across compartments, regardless of inbreeding, (3) heterosis played a much smaller role on microbial community diversity, composition, and function compared to tissue compartment or location, (4) in a small greenhouse experiment we found hybrid microbiomes failed to benefit inoculated MiniMaize.

## 5. Conclusions

Modern maize breeding is built on heterosis, and with many new biologicals coming to market, it is paramount to understand how these microbes interact with plant genetics. Our data show that although inbreeding has small and significant effects on taxa and function in maize microbial communities, these effects pale in comparison to the effect environment and tissue type have on community composition. With environment playing such a large role in shaping bacterial communities, investigating the extent that maize can recruit specific taxa/function would shed light on how much potential there is to use the plant itself to alter its microbiome. Further work is needed to examine how maize genetic diversity and the environment shape community function. Understanding how the microbiome interacts with maize genetic diversity will allow breeders and scientists to make better use of microbial communities for more sustainable crop production.

## Supporting information

Supplemental Figs and Files zip

Supplementary Materials: The following supporting information can be downloaded at:

Figure S1: AlphaDiversityBreakdown

Figure S2: InbredvsHybrid_PICRUST

Figure S3: MiniMaizeInoculation

Table S1: SharedTaxaUpset

Table S2: AlphaDiversityPvalues

Table S3: PERMANOVAresults

Table S4: MiniMaizeDriedMass

Dataset S1: Krona Plots

Dataset S2: Differential Abundance Tables

Dataset S3: PICRUST Tables

## Author Contributions

Conceptualization, J.W. and M.J.; methodology, C.S. and M.J.; field experimentation, C.S. performed the analysis and the MiniMaize inoculation experiment. M.J.; formal analysis, C.S.; writing—original draft preparation, C.S.; writing—review and editing, J.W.; visualization, C.S.; supervision, J.W.; project administration, J.W.; funding acquisition, J.W. All authors have read and agreed to the published version of the manuscript.

## Funding

This research was funded by the University of Georgia and the Foundation for Food and Agriculture Research (Grant #032).

## Data Availability Statement

All bioinformatics scripts and pipelines are available at: https://github.com/wallacelab/paper-schultz-microbiome-2023. The raw sequence data are available at the NCBI Sequence Read Archive under accession PRJNA924784.

## Acknowledgments

We would like to thank Holly Griffis for performing DNA extractions and library prep.

## Conflicts of Interest

The authors declare no conflict of interest. The funders had no role in the design of the study; in the collection, analyses, or interpretation of data; in the writing of the manuscript; or in the decision to publish the results.

## Notes

https://www.ncbi.nlm.nih.gov/bioproject/PRJNA924784

https://github.com/wallacelab/paper-schultz-microbiome-2023

### Competing Interest Statement

The authors have declared no competing interest.

