## Supplemental Figs and Files zip for "Effects of Inbreeding on Microbial Community Diversity of *Zea mays*": GeneticBackground-krona.html

Javascript must be enabled to view this page.

magnitude
magnitudeUnassigned

Blank
Hybrid
Inbred
Open\_Pollinated
Soil

5083558994152134917571918051

174913468117608775139

174913468117608775139

174913468117608775139

1739110869856697749

9059662813766093
1739110869856697749

242139210141

242139210141

112181355117

5811067137417629

123601750178986

123601750178986

123601750178986

123601750178986

92215494

92215494

92215494

92215494

4908657647350958916694417912

19340121376105474310087152

1550051026611

1550051026611

11512591

11512591

11512591

153453021556

153453021556

153453021556

2664192

2664192

2664192

141913

141913

141913

1734510024989888250366651

77649025962457111

6641839159521521
77649025962457111

8164942

1514242334644
231769288812433

83455546029

65638845585

166405855650
164685075048

1727862

1402122981261032462061

46454720832

46454720832

6095752739316621936

5755260711115801936

3449228282

329599950091552125

329599950091552125

10232585151101204401

43439592240888398
603581100746853

14051794

374238716326345

112108153

9686178427173163
309133217622393

532215627

9319126196

510519770

138544543610198338

138544543610198338
48114293283338

15148103118

15148103118

119723301891949

12580964483

2540

369222581004

331235245
369222581004

33612023759

101000232422

1011592

1011592

1089921733

1089921733

1089921733

581165451181752422361

20318251396432132

4561633115967

123501297

436641256

3179352421065

378147201042148102229

372145211035347962223

619968146

48868385686149144

120114276029625

120114276029625

180245117204731

16922461541404
180245117204731

2179

9188170691

61372158990

61372158990

6048238

6048238

6789964937310
1371659136559510

13778634

57683630188

4515420414

1070983569162701304

1070983569162701304
133557473997

760727849752126234

1771979145247658

21165

1401348343568776661329

1029791849504
1401348343568776661329

1299338553383871621329

198020627150765706490

588650649281339261

3382461249553451

1086857761159

216151714753235

2170794637

2170794637

128916550

128916550

25040452433805210
2282057145144192

221988982364118

221988982364118

718022025

718022025

718022025

718022025

13851394199284342229

427770460621716178
10381164783183441193

501641115266811

501641115266811

6681011

6681011

37316127664

37316127664

449649273383

449649273383

691269603607

691269603607

3472294161090136

3472294161090136
13890138037634

209139312305252

230587

230587

230587

230587

230587

16809308820295551948205805

1029214412017811741732936

64389768611342333935255

5183882831042473034442
2191222021047324924

662864

4003196189

7235417

46922942846055896

430447523448081601112

218119493727

140318881314

123865

188345351801239

4633774183

506700152

41277431

41277431

5237467114

11411031203408

216171620909

728642445521848210

1384789829917

6157191902

10823271103540101

83114957727290

51819441783647

51819441783647

232564131163

36013314713

36013314713

87222356141004451393

288936270933413

213396031864

256152133131

2274141969

917163421

58421420113914117380

3418119414772145
3741281856851592173

3346991538871128

94116859834519

90551728961235149
92641233281519187

289543228438

26461420356

223763603051

277321826211174202

277321826211174202

130265917581041178
277321826211174202

14755986313324

1381463

1381463

1381463

1381463

46611885210272309

466184989001769

466184989001769

100361212754

100361212754

2132466219944066

2132466219944066
1867821215715

27168498436961

195761612385877411

19837521635665511
195761612385877411

175924097502119

69210875665

69210875665

69210875665

89010474750922271009

79110148725521181008

2441414159316270
2441449165716574

356434

356434

3867377413021
3865475912118

191593

191593

371648738401368386

22339702152843139
371648738401368386

8965478016742

12728112

591647842310201

8938372

1211017538319466

1211017538319466
81251146157427

4076639216239

1752244613661
614813427

16457

63312519333
1135826710261

52716928

993012421091

1324158
992831511071

3582

3582

99116102471

99116102471

18912

18912

2512

2512

2512

2512

5627154226109925508613860

63125580638430

63125580638430

63125580638430
8115070738030

55105994

27351270113

49157

49157

49157

2721523161

2721523161

2721523161

872445

872445

872445

39839057569574237682290

2231121131
2287930334123544

176715126662089

382084

30430444226

4412479305863131

4412479305863131
4332294279163030

818526711

26823127125015

839339512113
22422196013

615117661

184308761292
920539832

9225478972

4512928

4512928

23715201368426149
2106643212

8844145011923

147973854275114

4161545697542751141

4161545697542751141

161152984

70214209146017200274
64439614331627

5326714102126583243

106309929566014

2161598311

1917477823586499611665
5411352889752396439

6101425013430306363

47414335707436451033

76869133

28556016316844127

42160163053272

42160163053272

691936936925
42160163053272

3668226116347

451730867496154

451730867496154

262144674

262144674

262144674

367935556582678170

142041132717

142041132717

353915155452651153
18131767433084

51346928749403
37179616044853

1416731270455

33322

2284340994244

441575337

275107483340022

46895745341676722

46895745341676722

11178010551303314

355327

355327

11174510021300614

11174510021300614

1561382365953270348

1561382365953270348

1561382365953270348
1561377965813226348

441444

8872453719792484456

8872453719792484456
3858274

455032301359

7702321116483464943

69738299753

92137711106952

92137711106952

92137711106952

92137711106952

92137711106952
1338177932

911039933602

1139273381815783181508

2276956429122

2276956429122

2276956429122

2276956429122

2276956429122

10940692585164080

10940692585164080

10938122413154380
10936352296140065

17711714315

17711714315

1318751
25717297

1268546

1268546

1030209931461662491306

588890663682661827

588890663682661827
4143522

26205289207
262693222694

6433624

342788150832090789
342783049482061789

3812010

131519

22071592025032
19526861133

1039828721429

15492233

206362225451221115

39116560932127
206362225451221115

22201110795

1451808154571764

1345630
893013

452617

3142257419
7767919

26245

21113460

223727542952043255

223727542952043255

118410724911134123
197554731741578191

1655316919610

3591659926

6344847

48426514925

23153890943962
1996346724527

445526417733

120178172

3190212262

831142

31079826

1311901408324109

1110371092301109
1110231082297104

141045

141045

215331623

215331623

215331623

3712659012966651290

3712659012966651290

1202296238776384

1202296238776384
3576959222522

57112282325561
57106978924261

533413

10474214

212214

212214

10446221

10446221

28176162341

16210289499833224

12910151492032834

38590359

913041200611114

652025552172

123630
331387839

33126429

33126429

434241230026

433568215019

433568215019
4352820462

394010417

6731507

66716

66716

61347

469764328124012

27423

27423

1226694102

1226694102

34947127642391

34947127642391

1322103428763269762

27571165994172248

51096177

51096177

2333172316076948

78984430

361357162525114
1553074311673948

6640045010420

6640045010420

409468582387

409468582387

1016073477

1016073477

1253476

1253476

3867623

3867623

3738352773876

3738352773876

118511745
3738352773876

357187

3632932469831

1563421

1563421

1563421

1563421

1563421

44163171026

44163171026

2772465

2772465

2772465

2339293961

2339293961

2339293961

237627768228

237627768228

237627768228

237627768228

237627768228

897204532345

806151551925

3124820
418104145

10656125

10656125

8019751475

8015338415

8015338415

44136

44136

910529842

910529842

910529842

910529842

459936174

459936174

459936174

459936174

459936174

892281755

811111401

811111401

811111401

811111401

8117354

8117354

8117354

8117354

53814

53814

3153
53814

22284

22284

16810338803073

54251230863

54251230863

54251230863

54251230863

114782650221

114782650221

114782650221

114782650221

271618425162475395296

404876453902482263

109132798327825

109132798327825

5234436510214
14248981009

389626725

5798361817611

295743744072204238

391042
182564120190691

12227110576

12227110576

4107439231061

286250341518

22972988416

5652053312

362186826

19630251693813135

19630251693813135

811848151348512

21891282316226

19988977316026

19988977316026

19988977316026

19988977316026

1923502

1923502

1923502

1923502

8234

8234

8234

8234

8234

209487411001127477

31022575

31022575

31022575

31022575

1802776987472479

1802776987472479

1802776987472479

28987010072637

252201205155

252201205155

252201205155

376698021087

376698021087

376698021087

11621817

11621817

11621817

11621817

11621817

11621817

32130820519010

416320

416320

416320
92918

32342

32342

922068813310

922068813310

92171417

92171417

92171417

1897411610

1897411610

1897411610

229615437

8307

8307

8307

8307

229532430

229532430

229532430

229532430

25238132786118437

25238132786118437

25238132786118437

25238132786118437
1339026

10713711386191

443823

10713271348168

1452309131096737
66317242

4351830014235

941711789814

217497

1461465

1461465

1461465

1461465

1461465

503518

503518

503518

503518

503518

503518

651623

651623

651623

651623

651623

651623

46918771574776

46918771574776

46918771574776

102952489

102952489

102952489

45915821326767

45915821326767

45915821326767

18411823218414565348

18411823218414565348

17691651917061450317

17691651917061450317

2726051168393

13899625123282453

35342763546164617
35342893565165717

131911

134232521945

1139
134232521945

2234121117

5748185

111214459

111214459

15624053

83448

83448

1482065

1482065

1482065

59113486190326

460836058514
59113486190326

293831112159

293831112159

232930660

232930660

311484433

311484433

4401376222806996262902

4401374392663995572902

4401374392663995572902

185725872316

185725872316
184804621883

92125433

289194431193948681459
606322382043874532855

132104603612

21610083652619661255

568747520167

831815103838262

125834774924159241

78826661615

47032114308157741

25191152690281

25191152690281

183143069

183143069

2721

2721

156140969

79132339

30457

474123
