## Supplemental Figs and Files zip for "Effects of Inbreeding on Microbial Community Diversity of *Zea mays*": GeneticBackground-Rhizos-krona.html

Javascript must be enabled to view this page.

magnitude
magnitudeUnassigned

Hybrid
Inbred
Open\_Pollinated

315273219715201

241319411817

241319411817

241319411817

214516251672

214516251672

214516251672

214516251672

268316145

200271119
268316145

684526

291143025613384

718859203290

201313951009

201313951009

683949
16910995

1017046

1017046

17231196777
18441286914

12190137

12190137

449339732068

11091256726

193652
11091256726

4188
8014315

24131

154214

9341030525
914955496

4419

167110

7647134
303283

461551

16021180688

16021180688

9613924

597724

3762

1221728524
457316185

19310267

332145178

23916594

285313140
201147114

20239

6414317

25345684

14721661
13520857

1284

1284

10624023

10624023

10624023

15291081570

802488275
15291081570

50644
24720467

8225

18911858

216248146

802813
1225430

422617

1428752

682552213

682552213

682552213

682552213

682552213

379135941369

127171111

127171111

203819

203819

203819

9811791

9811791

9811791

9161

9161

9161

24002405840

686603172

32020791
686603172

36138881

58

20302

20302

20302

234839160

3

3

1486582
234836160

5611672

306556

675936

675936

504236

1717

922

922

18312765

137

137

137

142210

372920

372920

1196935

1196935

1714638

853133

853133

853133

86155

86155

33035678

33035678
1781

281110

23229964

53383
151

52332

302232245

302232245

39811142

32410340

32410340

7482

7482

12641018418

936707305

31331382

24223639

717743

717743

561291197
623394223

6210326

6210326

328311113

328311113
283231112

45801

45801

22125364

22125364

22125364

22125364
948026

433823

433823

441114
8413515

402411

215

215

215

215

215

11457145475555

273498551842

128489369

2

2

126489369

126489369

180736331261

8520384

183512

3231003246
9282478664

8635

76160150

26201

191481187

595159

226178

1934117

236434114

23871173

1222631

1222631

11645142

302412214

281389200

212314

573754

573754

573754

1544

1544

1544

252713

252713

252713

5691247378

3701187314

3699817

5158

5344181

26212571

26212571

14537

1996064

1996064

1331463

1331463

1331463

1331463

719140233283

872244

2954

2954

2954

581740

581740

581740

986449215

986449215

988520249

988520249
38622472

411

16291

16

513737

515219139
18913028

32689111

508251243

508251243

508251243

15498152

13978125

13978125

152027

152027

1403190

1403190

1427138

1427138

1427138

649194526

649194526
35272

614167524

264721271379

144126

144126

4694
748751661

702657661

485379282
602741

14383

411314238

17715959

17715959

15111069
604221

916848

68034991

68034991

12211374

1019368

21206

16013364

14412148

161216

1109253

401044

401044

401044

401044

850250303

850250303

850250303
430138218

42011285

1532669430

1405596359

261817
14118

1279

1279

1006274188

841218147
1006274188

112264

533037

948867
756865

19202

19202

27921687
261310

9257

21716261
24417870

27169

1126571

222

222

813658
1106369

17239

17239

1242

1242

158

158

158

158

964704572

524755

524755

524755

524755

524755

20314616

20314616

14012710

14012710

14012710

63196

63196

63196

381921

381921

381921

381921

381921

480372325

1096177

1096177

322209203

116106104
322209203

1253476

1253476

816923

816923

4910245

4910245

4910245
316345

1839

1104

1104

16
1104

44

44

190110151

190110151

33136

33136

33136

534820
15797145

10449125

10449125

351621

351621

351621

351621

351621

351621

5541956212

373526143

373526143

15820154

15820154

21532589

12520246
21532589

9012343

181143069

181143069

2521

2521

156140969

474123

79132339

30457

14961127

14248114

14248114

14248114

14248114

14248114

117

117

117

117

117

726

726

726
224

52

52

628557411

628557411

7445145

6726115

24

6723111

1

71930

518445244

268283119
518445244

52714

835045
492932

342113

482310

10188

10188

1044448

1044448

323

323

323

334422

6275

6275

271717

271717

271717

307820681223

269615481064

435341125

435341125

1905782

24528443

22611207939

475498189

231042
1771701739

1416136

1416136

1002367301

337135308

337135308

26812852

15811

15811

3152

3152

3152

3152

3152

379505157

379505157

159164155

159164155

159164155

2203412

2203412

2203412

302816

302816

302816

302816

302816

302816

4257

4257

4257

4257

4257

4959

4959

4959

4959

4959

964453517

964453517

535186256

854524
535186256

34468199

34468199

1067333

1067333

32996181

29
32996181

18228117

1194259

1194259

26175

5213053

481035

481035

481035

42748

42748

484127

484127

484127
352516

131611
