## Supplemental Figs and Files zip for "Effects of Inbreeding on Microbial Community Diversity of *Zea mays*": GeneticBackground-Roots-krona.html

Javascript must be enabled to view this page.

magnitude
magnitudeUnassigned

Hybrid
Inbred
Open\_Pollinated

36023444515151

2103268

2103268

2103268

43

43

43

43

2063265

2063265
183305

23215

35813441255143

295320028

5666

2410

2410

2410

2410

3256

3256

3256

3256

103522

3582

3582

3582

3582

6844

6844

6844

6844

228116514

1762614

1101894
1762594

5754

5754

916

916

2

21051390

21051390

21051390
15454

339145

16121191

302362

2029

2029

2029

2029

28272

2617
259

27

27

212

11

11

102

102

141111

141111

145

145

145

127106

127106

127106

5271

5271

5271
3047

2224

2224

1815

1815

1815

1815

1815

28589746

28589746

1043

517

526

526

20127042

20127042
1388936

1146

1146

18153

159

2

2

1913

1913

614812

35762

35762

25

25

24400

24400

131032

1390

3
1390

1387

132

132

662834308

662834308

258387

258387

204253

54134
54131

3

9413613

9413613
930

22

22

313113

5273

225201295

225201295
209174285

81110

81110

816

816

85110

81103

81103

81103

47

47

1759922

1759922

1759922

145

123
145

22

1307117

1086517

8

146

31235

31235

216

216

216

216

216

216

16698298432136

703020062865

50631

50631

50631

34523723

36

36

34223123

34223123

2

2

2

781996121

985138

58578

40456

683483113

19523586

48824827

5

5

5

5

1520

1520

1520

443271111

443271111

443271111

538917907610

3858

41434

548

548

36386

1875

6774

1025434735
474816923526

2095942

26322533372

40774

1523

1811851

92

5467485113

813

47734384

292815

24324629

2056940

70667321912

420187109

420187109

420187109

41563970452

102107

102107

102107

1066730

96768422
19801473137

266100112

7476893

2730
38395725

35692725

462211
33930968

16315847

13012910

24041112

84136
1923442

1082082

486710

27882

27882

1922767
1834

75116

821265

172

787282171

787282171

1207329

1207329

1207329

1795032

1795032

42

42

42

42

856213775

19335
856213775

1483131

567195315

12212024

922626189

794571171
663318171

87233

4420

1285518

1285518

49178

20103

20103

20103

2975

2975

2975

29121313

29121313

22516013
29121313

6653

49135

49135

2033

2033

2033

26022460359

1131132

1689

1689

515
97242

23152

23152

234

234

104

104

104

104

24792343357

47893810
4628949

16441

16441

18901238324

17431088275
18901238324

1527

9614042

368

57907
9015121

264114

720

1482
21162

78

78

555531

555531

555531

555531
3910

503507

1

502507

1314

1

1214

572448041217

96749

96749

53

53

53

80379

80379

80379

58

58

58

626

626

626

1844975287

1122477211

20710453

59919

59919

1489534

32014635
915373158

595227123

595227123

4941

4941

4941

4941

67345776

25133

25133

64844473
44633556

572117

572117

14588

14588

37843755921

22815624

2210

2210

714320

714320

278

278

42

236

7471

1151

1151

23193

23193

682913

622713

562513

62

62

62

18864109

18864109

295

293
295

2

11016423

2951

2951

8151
8111323

739822

8031844

803184

803184

803184

4

4

1375213918

354211
1375213918

11872019
121620453

29263

47
124524

120454

447517633

436488633
447517633

36

823

1271706

1271706
791232

32394

168

40932791

37731691

37731691

2411

2411

8

8

741241661246

88175543

88175543

88632
63482

2515

2515

79369241
73766535

56276

56276

1594404216

1594404216

1594404216

1594404216

1594404216

49373007987

51127451

1303518
51127451

129

2735
50217

22

21162

122
92

3

30720726

28211429555

28211429555

94732

48212

4652

29512019
54424963

5635

1939444

21831107490
1772944403

24510055

1666332

39827266

34919466
35119666

22

22

4776

4776

4776

12071032315

227
12071032315

908614270
939674270

2249

911

2616
443411

181811

85221
20231734

1179634

15

15

15

15

15

6283

6283

6283

6283

6283

6283

302

302

302

302

302

302

9111916

346114

346114

346114
72714

2734

2734

1030

819

819

819

819

211

211

211

211

47282

47282

47262

47262

47262

2

2

2

354

354

354

354

354

2633

2633

2633

2633

2633

1064682140

705435138

26416243

26416243

17115
49325

3221

21513038

44127395

16
42823890

945830

31818060

2287023

9011037

13355

582

582

582

582

582

354239

39149

39149

39149

39149

4245

4241

4241

4241

4

4

4

27345

27345

27345

2072

2072

2072

2072

2072

2072
