## Supplemental Figs and Files zip for "Effects of Inbreeding on Microbial Community Diversity of *Zea mays*": GeneticBackground-Stalks-krona.html

Javascript must be enabled to view this page.

magnitude
magnitudeUnassigned

Hybrid
Inbred
Open\_Pollinated

722805943028188

719615908328188

1441165

1441165

1441165

1441165

1441165

97171

9735

9732

9732
66

3

3

2832

2832

3

3

3

3

136

136

136

136

136

216

216

216

216

2

2

16

193

193

193

193

193

462464599722403

157591168620936

49

49

21

21

21

21

123446612872

123446612872

123446612872

23

23

19

4

1018096306868

4058923
858469485607

49

452659765607

4

4

449

449

52546
525503

3

4

210
152416581258

131416581258

1611

1611

14

14

14

432215111194

432215111194

432215111194

7

7

7

7

22

22

2
22

2

30485342721465

1123115813137

100311212154

100311212154

1200369283

1200369283

78

78
62

16

18435169551165

31

4915233552
18009166131155

2121

65

45

1712026

120978281026

8

7811123651

35

35426910

35426910

72

7

7

8191426163

1074

24

105

7121426159

6321383159

8

8

8035

8035

2392

2392

7

2

2

2

2

2

2

32

28148

94

94

94

27244

176

176

17

16

1

6

847

847

847

7

7

16431

16431

31

164

18642

2

2

2

2

16642

16642

342

302

2

2

1630

1630

37589

37589

18537

18537

1323

5514

19

19

19

19

43

43

43

43

9

9

9

743256551045

16151053

1628253

1618353

69

69

166453

50

99

35

35

35

35

145193

145179
143175

24

24

14

14

14

12

12

12

12

12

72595145992

114

114

109

5

33612021759

33612021759

33612021759

15

15

15

204664765

196333247
197463547

11303

721218

4454

4454

4454

6715866

4317
6715866

2414166

42142929

201465

201465

8429124

8429124

13692

13692

3145

73

69

4

372

372

1287155560

1266150260

1266150260

1920

26

26

27

152213

152213

151713

5

1118

1118

1118

8

8

8

111

111

111

1759362264733

1759362264733

716428852168

716428852168

59

651525272168

590358

81703173

81703173
19

120

2

81562

173

173

968125282392

216144

216144

946525142388

946525142388

667110

667110

66716

66716

94

319347

319347

319347

319347

319347
42

315314

31

31
