## Supplementary figures and images for "Effects of Inbreeding on Microbial Community Diversity of *Zea mays*"

### FigS1_Alpha.png

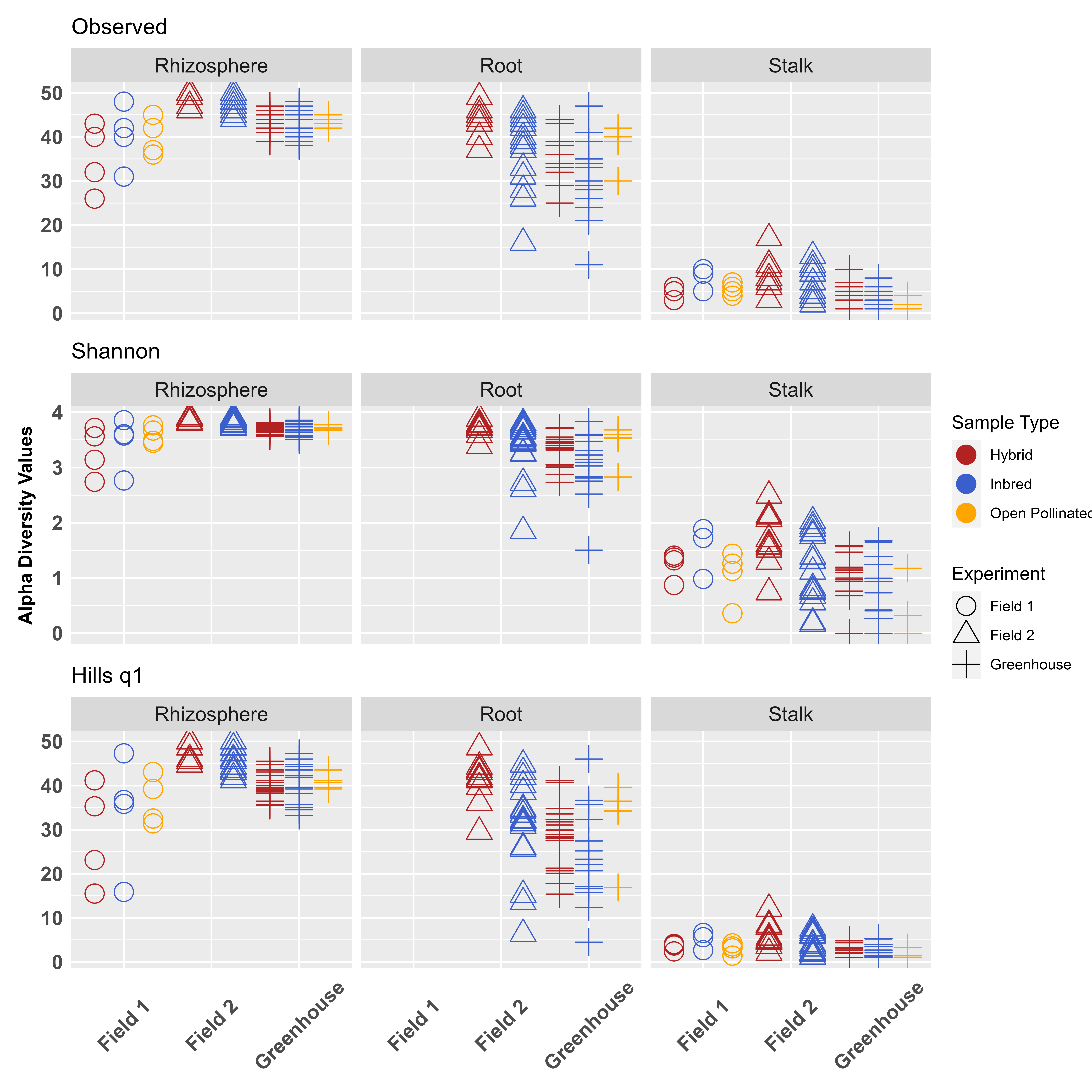

### FigS2_IvHpicrust.png

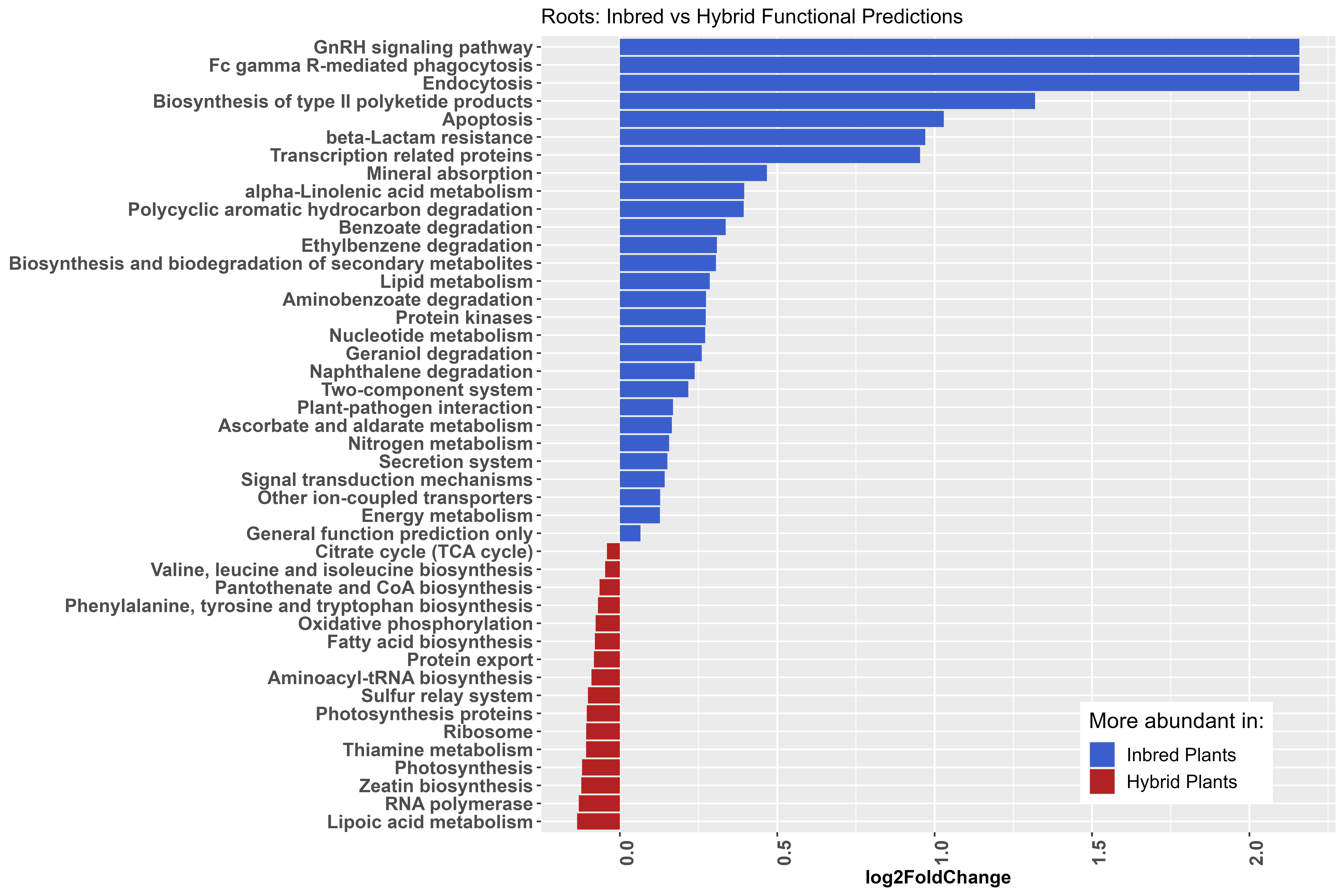

### FigS3_MMinoc.png

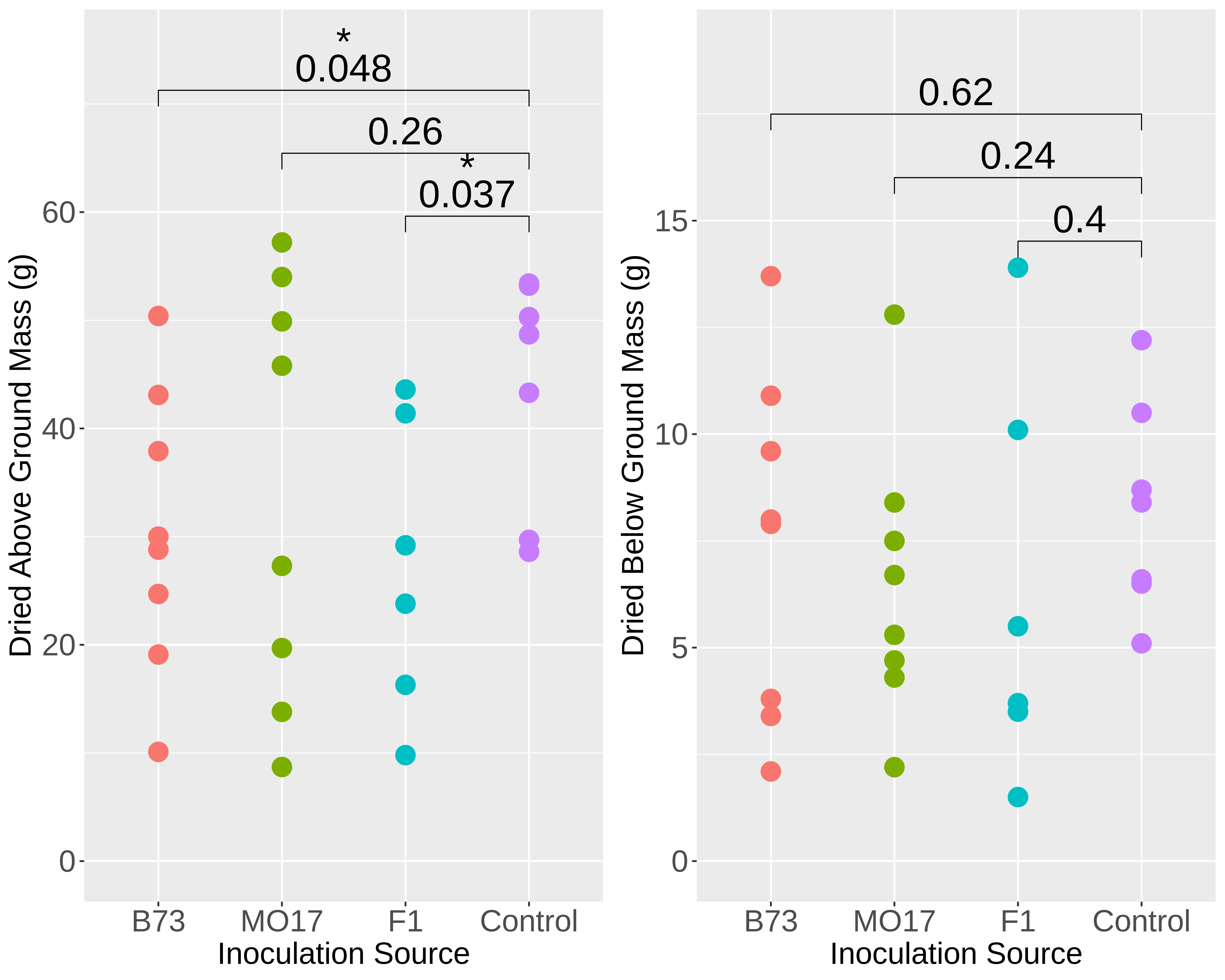

### Roots_Hybrid_Krona.png

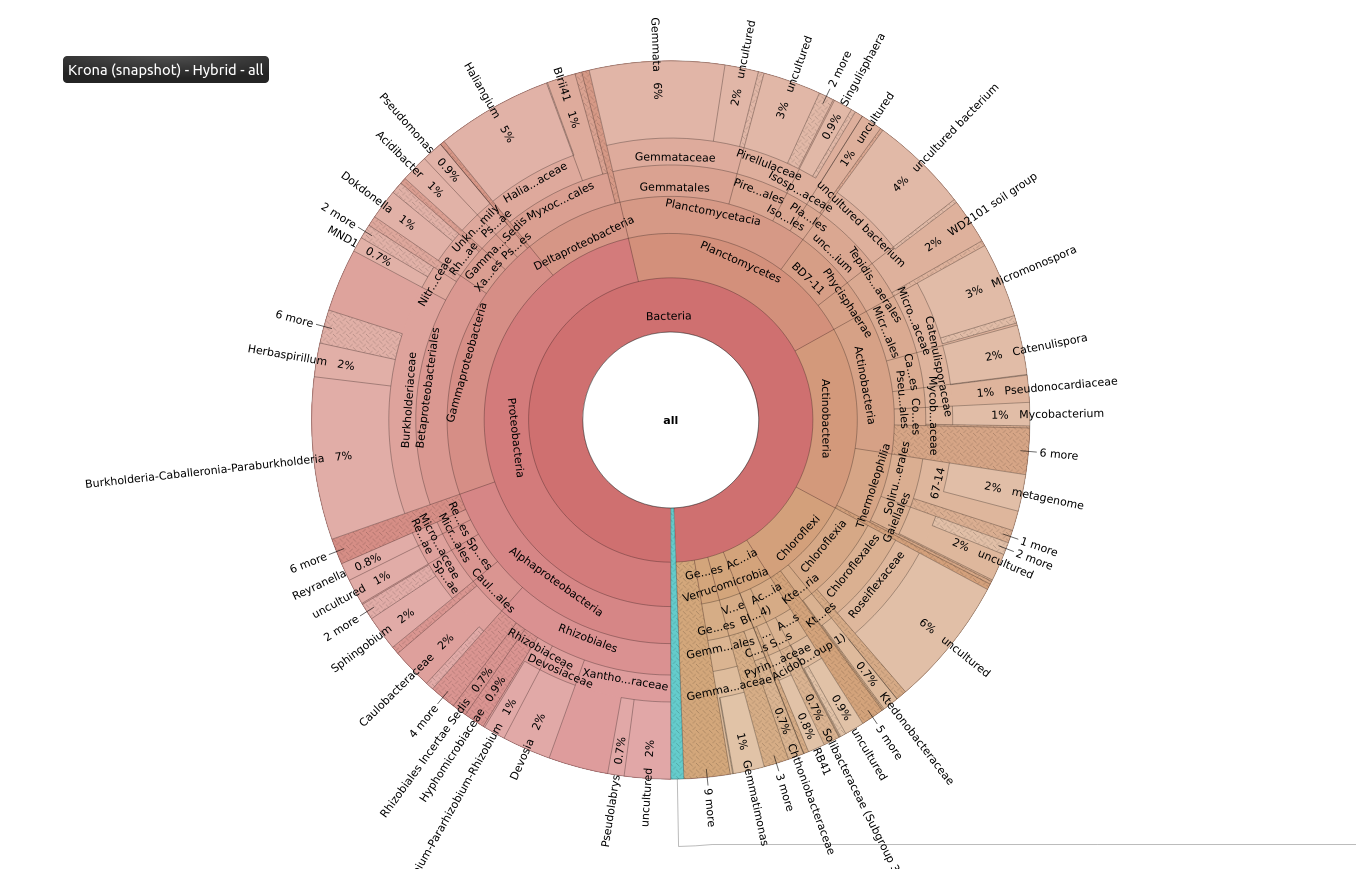

### Roots_Inbred_Krona.png

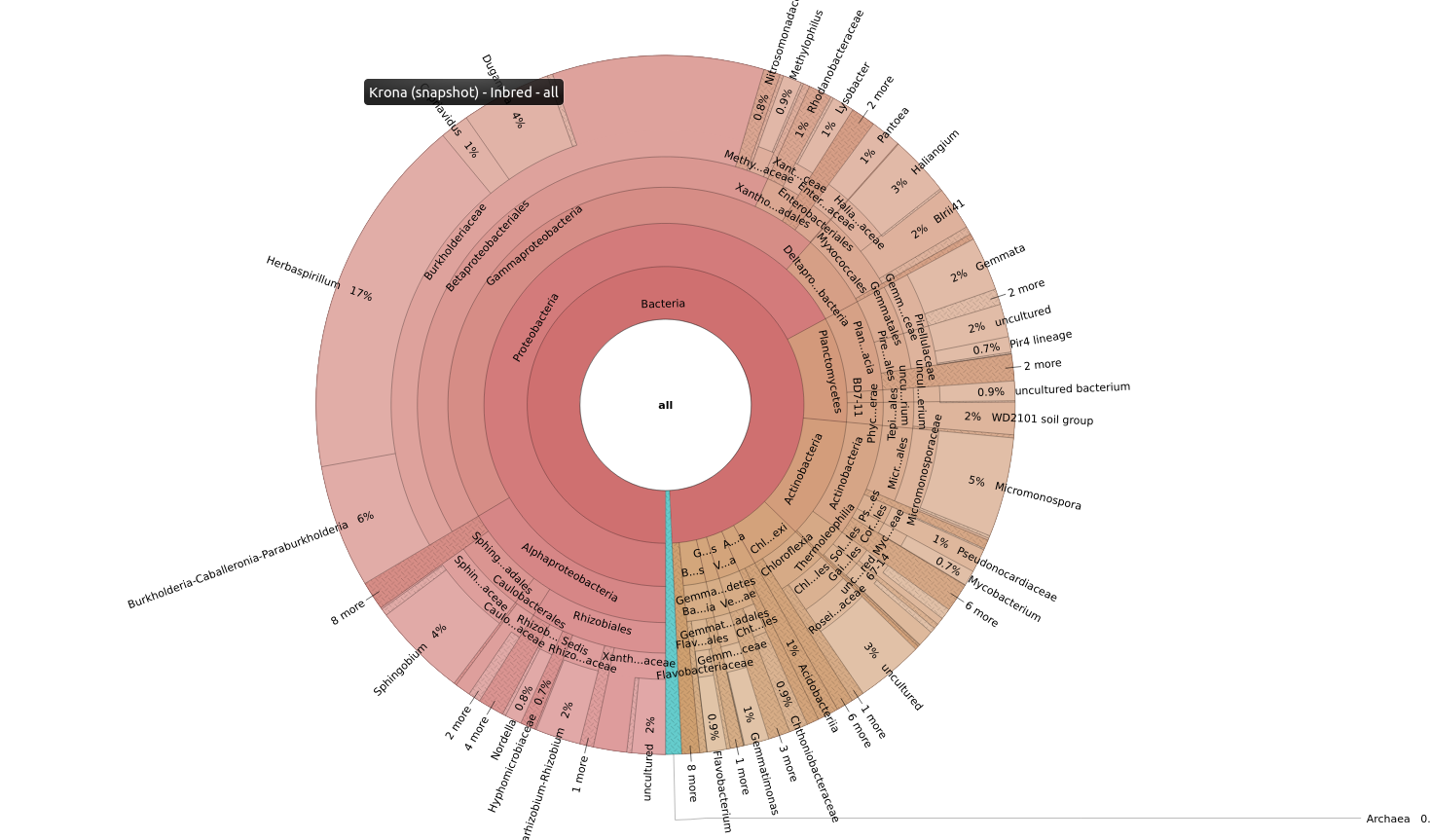

### Roots_OpenPol_Krona.png

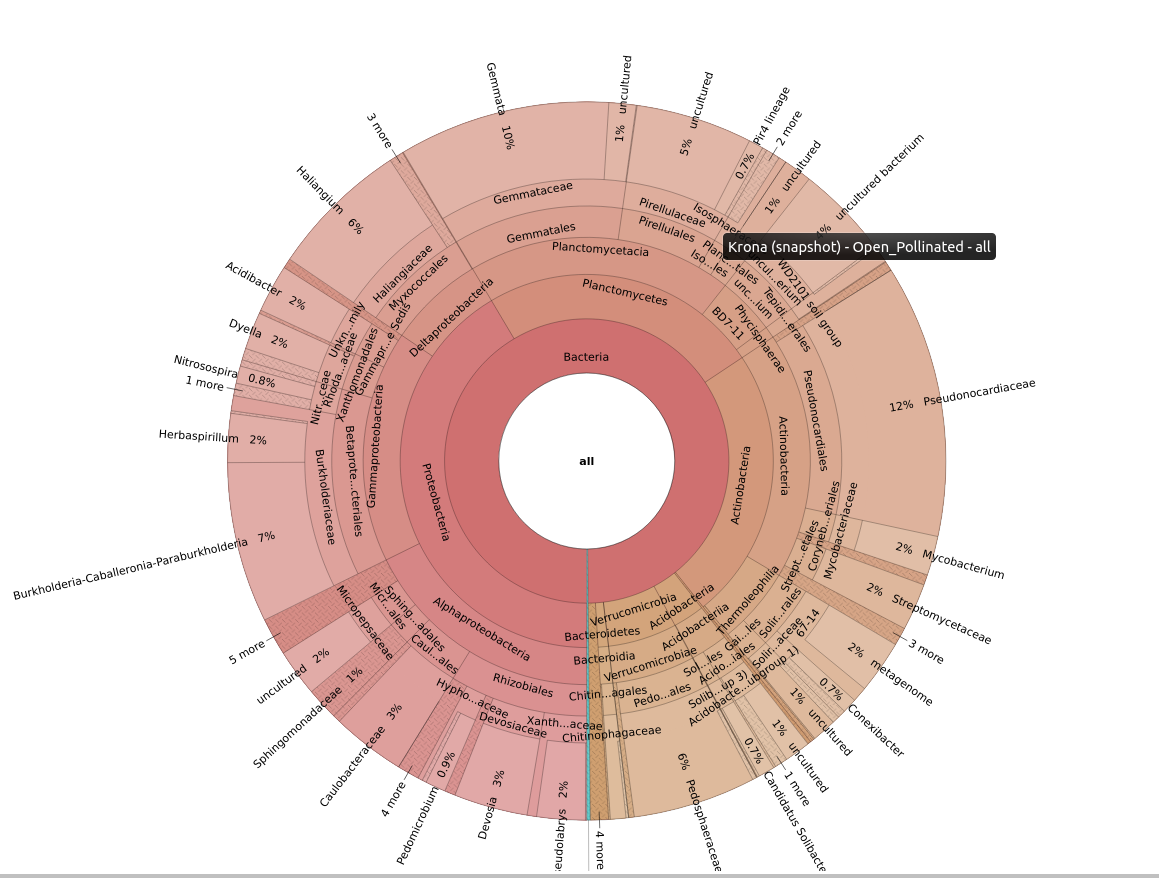
